# Using Twitter To Generate Signals For The Enhancement Of Syndromic Surveillance Systems: Semi-Supervised Classification For Relevance Filtering in Syndromic Surveillance

**DOI:** 10.1101/511071

**Authors:** Oduwa Edo-Osagie, Gillian Smith, Iain Lake, Obaghe Edeghere, Beatriz De La Iglesia

**Affiliations:** School of Computing Science, University of East Anglia, Norwich, Norfolk, United Kingdom; Real-time Syndromic Surveillance Team, National Infection Service, Public Health England, Birmingham, United Kingdom; Epidemiology West Midlands, Field Service, National Infection Service, Public Health England, Birmingham, United Kingdom; School of Environmental Sciences, University of East Anglia, Norwich, Norfolk, United Kingdom

## Abstract

We investigate the use of Twitter data to deliver signals for syndromic surveillance in order to assess its ability to augment existing syndromic surveillance efforts and give a better understanding of symptomatic people who do not seek health care advice directly. We focus on a specific syndrome - asthma/difficulty breathing. We outline data collection using the Twitter streaming API as well as analysis and pre-processing of the collected data. Even with keyword-based data collection, many of the tweets collected are not be relevant because they represent chatter, or talk of awareness instead of suffering a particular condition. In light of this, we set out to identify relevant tweets to collect a strong and reliable signal. For this, we investigate text classification techniques, and in particular we focus on semi-supervised classification techniques since they enable us to use more of the Twitter data collected without needing to label it all. In this paper, propose a semi-supervised approach to symptomatic tweet classification and relevance filtering. We also propose the use of emojis and other special features capturing the tweet’s tone to improve the classification performance. Our results show that negative emojis and those that denote laughter provide the best classification performance in conjunction with a simple bag of words approach. We obtain good performance on classifying symptomatic tweets with both supervised and semi-supervised algorithms and found that the proposed semi-supervised algorithms preserve more of the relevant tweets and may be advantegeous in the context of a weak signal. Finally, we found some correlation (*r* = 0.414, *p* = 0.0004) between the Twitter signal generated with the semi-supervised system and data from consultations for related health conditions.

## 1 Introduction

Surveillance, described by the World Health Organisation (WHO) as “the cornerstone of public health security” [1], is aimed at the detection of elevated disease and death rates, implementation of control measures and reporting to the WHO of any event that may constitute a public health emergency or international concern. Disease surveillance systems often rely on laboratory reports. More recently some countries such as the UK and USA have implemented a novel approach called “syndromic surveillance”, which uses pre-diagnosis data and statistical algorithms to detect health events earlier than traditional surveillance [2]. Syndromic surveillance can be described as a real-time (or near real-time) collection, analysis, interpretation, and dissemination of health-related data to enable the early identification of the impact (or absence of impact) of potential human or veterinary public health threats that require effective public health action [3]. For example, they use emergency department attendances or general practitioner (GP, family doctor) consultations to track specific syndromes like influenza-like illnesses (ILI). The expansion in digital technology and increasing access to online user-generated content like Twitter has provided another potential source of health data for syndromic surveillance purposes. Expanding access to communications and technology makes it increasingly feasible to implement syndromic surveillance systems in low and middle income countries (LMIC) too and some early examples in Indonesia and Peru have indicated reasons for optimism [2].

The use of data from microblogging sites such as Twitter data for disease surveillance has been gaining momentum (e.g. [4–8]). This may not only complement existing surveillance systems but may support more accurate monitoring of disease activity in sub-groups of the population that do not routinely seek medical help via existing healthcare services. The real-time and streaming nature of Twitter data could provide a time advantage for syndromic surveillance activities aimed at early detection of disease outbreaks. In addition to this, the low cost of utilisation of this data means that in LMIC where access to medical services may be restricted but where the of use digital technology and social media is becoming more common, such data may support the development of cost-effective and sustainable disease surveillance systems.

It is in this light that we develop our work. Our aim is to establish the utility of social media data and specifically Twitter data for syndromic surveillance.

Our first objective is to extract a reliable signal from the Twitter stream for different syndromes and health conditions of interest. To achieve this, we must be able to effectively identify and extract tweets expressing discomfort or concern related to a syndrome of interest and reflecting current events. Such symptomatic tweets are considered “relevant” for our purpose of syndromic surveillance. In this paper, we look at at asthma/difficulty breathing as our syndrome of interest which has received less attention in studies using social media data than other syndromes (e.g. ILI). Simply using the keyword “asthma” or other asthma-related keywords associates with tweets such as “*oh I used to have asthma but I managed to control it with will power*” or “*Does your asthma get worse when you exercise?* “which we consider as not relevant. On the other hand tweets such as “*having an asthma attack atm*” or “*why is my asthma so bad today?* “express a person currently affected and we would like to consider them as relevant. Hence the problem becomes one of text classification. Other authors [8, 9] have already identified that much of the data captured on Twitter represents chatter, concern or awareness instead of actual infection or suffering from symptoms of a disease, talk of past events or a reflection on news content, and is therefore irrelevant as a signal for syndromic surveillance. Such irrelevant content may greatly magnify the signal and lead to incorrect results and over-estimation [10]. Once the signal has been extracted, we then compare it to real world syndromic surveillance data to understand how well Twitter works for monitoring our syndromes.

When extracting the Twitter signal, we focus on two novel aspects of tweet classification. Firstly, we investigate emojis in tweet classification, and show their worth in a syndromic surveillance context. While there is published literature making use of emoticons in text classification, there are few studies describing the use of emojis. Secondly,we compare both supervised and semi-supervised approaches to text classification. We consider semi-supervised methods because they enable us to use unlabelled data, thereby reducing the initial labelling effort require to build a classifier. Finally, we compare the signal we extracted using our methods to syndromic surveillance data from Public Health England (PHE) to investigate the utility of Twitter for the syndromic surveillance of asthma/difficulty breathing.

## 2 Related Work

In a survey carried out in 2015, Charles-Smith et al. [5] identified 33 articles that reported on the integration of social media into disease surveillance with varying degrees of success. However, they reported that there is still a lack of application in practice despite the potential identified by various studies. Many studies are retrospective as it is relatively easy to predict a disease post outbreak but practical application would need to be prospective. Uses of social media data vary from global models of disease [11] to the prediction of an individual’s health and when they may fall ill [12].

The most commonly studied disease is influenza or ILI [13]. Ginsberg et al. [7] put forward an approach for estimating influenza trends using the relative frequency of certain Google search terms as an indicator for physician visits related to influenza-like symptoms. They found that there was a correlation between the volume of specific Google searches related to ILI and the recorded ILI physician visits reported by CDC [7]. De Quincey and Kostkova [6] introduced the potential of Twitter in detecting influenza outbreaks. They posited that the amount of real-time information present on Twitter, either with regards to users reporting their own illness, the illness of others or reporting confirmed outbreaks from the media, is both rich and highly accessible. Achrekar et al. [4] also investigated the use of Twitter for detecting and predicting seasonal influenza outbreaks and observed that Twitter data is highly correlated with the ILI rates across different regions within USA. They concluded that Twitter data can act as supplementary indicator to gauge influenza within a population and could be useful in discovering influenza trends ahead of CDC.

In this study, our objective is to collect relevant tweets for our given syndrome. We proceed to the initial data collection by using a set of possibly related keywords. However, we notice that a majority of tweets are not relevant as they do not express the required sentiment (i.e. a person suffering from the particular ailment at the current time). We then view this as a text (or tweet) classification problem and build models to filter relevant tweets. Several papers have looked at the tweet classification problem using supervised learning for different applications. Sriram et al. [14] classified tweets to a predefined set of generic classes such as news, events, opinions, deals, and private messages, based on information on the tweets’ authors and domain specific features extracted from tweets such as the presence of abbreviated words. Dilrukshi et al. [15] applied a Support Vector Machine (SVM) to classify tweets to different news categories. The most relevant work in the context of tweet classification is that of Dredze and his colleagues [8, 9] as they used Twitter data to investigate influenza surveillance. They argue that for accurate social media surveillance it is essential to be able to distinguish between tweets that report infection and those that express concern or awareness. One problem with these approaches is that they rely on having a set of labelled data for learning, i.e. a sufficient set of tweets must first be labelled as say relevant/irrelevant for the learning to take place. Such labelling can be very time consuming so it often means that researchers do not use all of the data available but instead use a subset of labelled data to develop their classifiers. Since the syndromes/events we wish to study may not be mentioned frequently in a Twitter feed, we wish to use as many tweets as possible to build our models. To this effect semi-supervised classification approaches try to produce models using a small set of labelled data but also taking into account the larger set of unlabelled data so we investigate them next.

Zhang et al. [16] investigated the semi-supervised classification of tweets for organization name disambiguation, a problem previously tackled with a supervised approach by Yerva et al. [17]. Zhang et al. compared Label Propagation and Transductive Support Vector Machines (TSVMs): both methods utilise unlabelled data in the classifier. A number of papers have looked at using semi-supervised learning for sentiment analysis, and in particular self-training [18, 19]. Baugh [20] proposed a hierarchical classification system with self-training incorporated into it where his goal was to classify tweets as *positive*, or *neutral*. Liu et al. [21] proposed a semi-supervised framework for sentiment classification in tweets that was based on co-training. They converted tweets into two kinds of distinct features - textual and non-textual. Two Random Forest (RF) classifiers were trained with the same labelled data but one with textual features and the other with non-textual features. Johnson et al. [22] proposed a general semi-supervised framework for document classification using Convolutional Neural Networks. Lee et al. [23] applied this framework to the classification of tweets as being related to adverse drug effects or not.

In this paper, we build classification models for tweets based on the relevance in the context of a specific syndrome/event. As part of our investigation into feature representation and feature selection for text, which is an important part of text classification, we experiment with different types of features, taking into consideration suggestions from previous work. We also consider emojis in tweet classification, and show their worth for tweet classification in a syndromic surveillance context. We compare both supervised and semi-supervised approaches to text classification in order to understand if and how we can utilize more of the data that we collect.

## 3 Methods

We discuss the data collection, pre-processing and analysis of tweets in order to extract a relevant signal for a given syndrome. We narrow our efforts to asthma and air pollution incidents in this paper.

### 3.1 Data Collection and Pre-processing

Tweets were collected over multiple periods to account for seasonality in asthma activity and to have a higher chance of an air pollution event being observed. Different periods also enable us to monitor changes in the use of Twitter as well as in the language used on Twitter over time. We started with an Autumn period (September 2015 to November 2015), followed by a summer period (June 2016 to August 2016) and a winter through to mid-summer period (January 2017 to July 2017).

Tweets were collected using the official Twitter streaming Application Programmer’s Interface (API). The Twitter streaming API provides a subset of the Twitter stream free of charge. The whole stream can be accessed on a commercial basis. Studies have estimated that using the Twitter streaming API, users can expect to receive anywhere from 1% of the tweets to 40% of tweets in near real-time [24]. The streaming API has a number of parameters that can be used to restrict the Tweets obtained. We extracted Tweets in the English language with specific terms that may be relevant to a particular syndrome. For this, in conjunction with experts from Public Health England (PHE), we created a set of terms that may be connected to the specific syndrome under scrutiny, in this case asthma and difficulty breathing. We then expanded on this initial list using various synonyms from regular thesauri as well as from the urban dictionary (https://www.urbandictionary.com) as that may capture some of the more colloquial language used in Twitter. Examples of our keywords are “asthma”, “wheezing”, “couldn’t breathe” etc. A full list of terms used is provided in the appendix.

We collected 10 million tweets obtained over the three collection periods. The general characteristics of the collected tweets are reported in table 1. The anatomy of a Tweet is presented in the Status Map in Fig 1. There are a number of attributes that are associated with a Tweet and would be available to our analysis. We did not consider all the available Tweet attributes useful for our experiments so we collected those that could help us in our task. More specifically, we collected “Tweet_Id”, “text”, “created_at”, “user_id”, “source” as well as information that may help us establish location such as “coordinates”, “time_zone” and “place.country”. We stored the collected Tweets using MongoDB, which is an open source no-SQL database whose associative document-store architecture is well suited to the easy storage of the JSON Twitter responses.

**Table 1.** Information on the data corpus collected before cleaning

|  | Counts |
| --- | --- |
| Tweets | 10,702,063 |
| URLs | 2,225,155 |
| Hashtags | 177,506 |
| Emojis | 3,103,598 |
| Number of unique users | 5,861,247 |
| Number of tweets per user | 4.1 |

**Fig 1.**
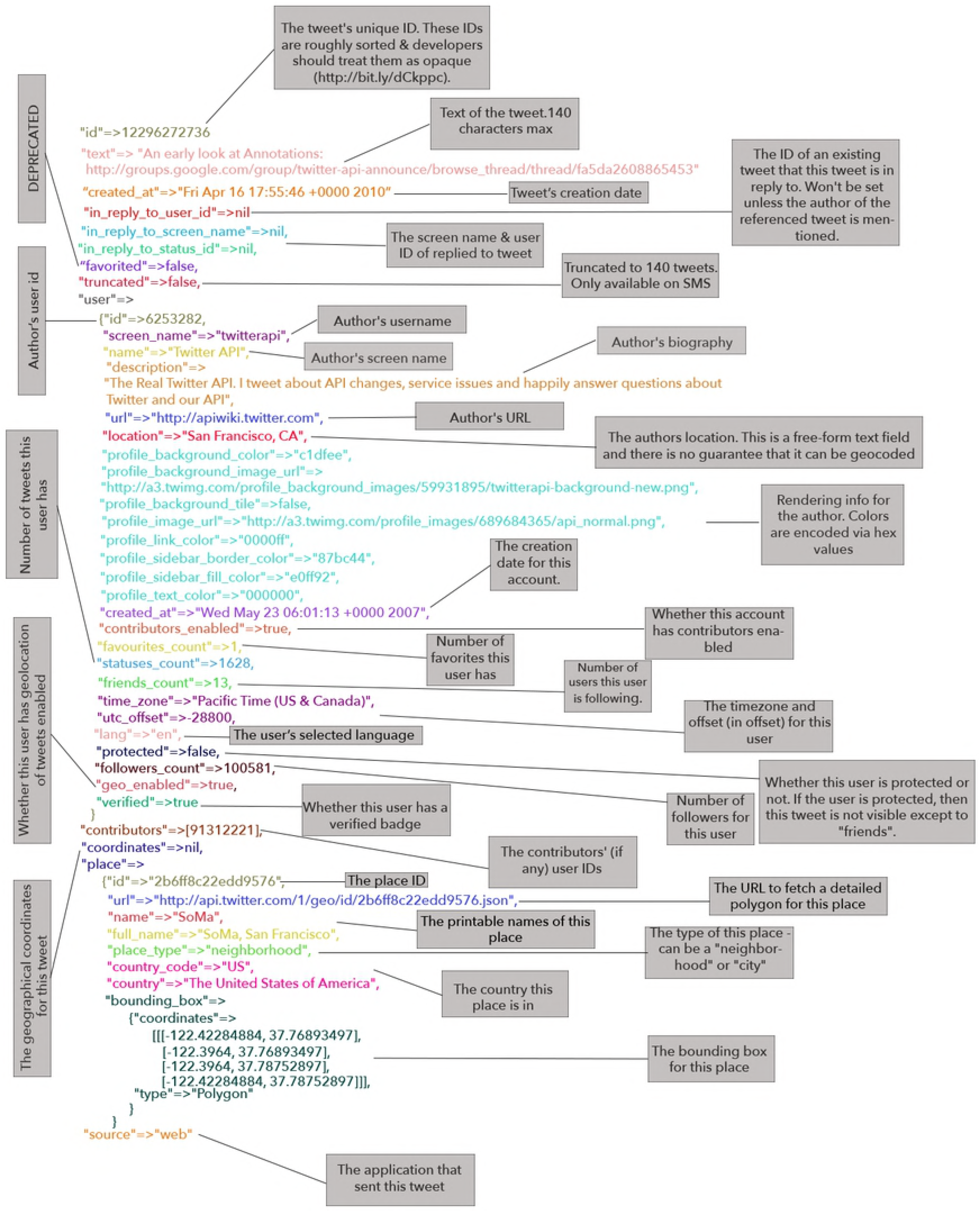
Map of a Tweet from the Twitter API

#### 3.1.1 Location Filtering

Because the aim of this project is to assess the utility of Twitter data for syndromic surveillance systems in England, we would like to exclude tweets originating from outside England. Doing this will give a realistic signal, however, inferring the location of Twitter users is notoriously difficult. Fewer than 14% of Twitter users disclose city-level information for their accounts and some of those may be false or fictitious locations [15]. Less than 0.5% turn on the location function which would give accurate coordinate information from mobile devices. *time_zone, coordinates* and *place* attributes, which we collected, can help in the geolocation of a tweet but are not always present or even correct as is shown in table 2. The most reliable at the time of tweeting, coordinates, is only present in a very small percentage of tweets.

**Table 2.** Availability of geolocation attribute in collected Twitter Data

| Data Collection Period | Percentage of Tweets Containing attributes |  |  |
| --- | --- | --- | --- |
|  | Coordinates | Timezone | Place |
| September 23, 2015 - November 30, 2015 | 0.30% | 57.90% | 2.17% |
| June 15, 2016 - August 30, 2016 | 0.29% | 61.12% | 2.10% |
| January 27, 2017 - July 31, 2017 | 0.21% | 59.21% | 1.61% |

For building a relevance classifier, accurate location is of relative importance. In this work, we are not overly concerned with accurate location filtering. For the purpose of symptomatic tweet classification for relevance filtering, location is of no importance. We collect tweets from the whole of the UK. We employ all three geolocation fields, filtering out tweets that do not have a UK timezone, a place in the UK or coordinates in the UK. We acknowledge that the location filtering is not entirely accurate and may have a disruptive effect when we compare our signal with public health data collected within England. However, we leave the task of improving on location filtering for future work where we will extend our signal comparisons to include longer periods of time and other syndromes.

#### 3.1.2 Cleaning the Data

The Twitter dataset contained retweets (sometimes abbreviated to RT) which are the re-posting of a tweet; other duplicate tweets not marked as RT but containing exactly the same text with different URLs appended; and users tweeting multiple times on the same day. We removed all duplication from the final dataset in order to minimise the detection of false signals

In addition, we removed URLs, which are often associated with news items and blogs, and replaced them with the token “<URL>“. This helped with identification of duplication but also identification of “bot” posting and news items. A “bot” is the term used when a computer program interacts with web services appearing as a user. Tweets from bots, news and web blogs are not relevant to syndromic surveillance so we developed algorithms to identify them and remove them. An overview of the data after cleaning, showing a considerable reduction in volume, is shown in table 3.

**Table 3.** Information on the data corpus collected after cleaning

|  | Counts |
| --- | --- |
| Tweets | 127,145 |
| URLs | 147,102 |
| Hashtags | 23,189 |
| Emojis | 36,872 |
| Number of unique users | 115,583 |
| Number of tweets per user | 5.3 |

#### 3.1.3 Labelling

3,500 tweets from the first data collection period were manually labelled as “relevant” or “not relevant”. A tweet was labelled as relevant if it announced or hinted at an individual displaying symptoms pertaining to the syndrome of choice. The labelling was done by three volunteers. A first person initially labelled the tweets. This took approximately 1 hour per 1,000 tweets. A second person checked the labels and flagged up any tweets with labels that they did not agree with. These flagged tweets were then sent to the third person who made the decision on which label to use. 23% of the labelled tweets were labelled as “relevant” while 77% were labelled as “irrelevant”. A second set of 2,000 tweets, selected at random, were later labelled following the same procedure from the last data collection period. 32% of these tweets were labelled as relevant and 68% were labelled as irrelevant.

#### 3.1.4 Basic text classification features

Although it is possible to use any sequence of letters or language tokens to represent text, words have been used successfully in language modelling and speech recognition [25]. Words are identified after a process of tokenisation and can then be used to represent a document by their presence or absence without trying to retain any information on the ordering of words, their frequency or their relationship to one another. That approach is called “bag of words” and despite its relative simplicity can work well in many text mining scenarios. It is also possible, with the bag of words model, to use weighting schemes such as tf-idf (Term Frequency-Inverse Document Frequency) [26] and they may perform better than a boolean representation. However, some authors [8, 27] have argued that more complex features will dramatically decrease the feature space while leading to better classification performance. We acknowledge that deep learned word vectors are an effective avenue for text feature representation. However, the training and deployment of deep learning systems can be intensive and require considerable hardware resources. Instead of going down the deep learning route, we look towards building the cheapest and simplest system possible with little or no compromise on effectiveness. We believe this will make it easier for low and middle income countries (LMIC) to incorporate such systems at whatever scale.

Classification of tweets may be challenging as tweets are very short and in our scenario, the classes may share common vocabularies. That is, both relevant and irrelevant tweets could contain the same words. Twitter has specific language and styles of communication that people use. In particular, we found that *emojis and emoticons* are promising additional tokens that we could exploit in classification:

- An emoticon is a pictorial representation of a facial expression using punctuation marks, numbers and letters, usually written to express a person’s feelings or mood.:-) is an example of an emoticon.
- Emojis on the other hand are miniature graphics of various objects and concepts including facial expressions. 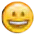 is an example of an emoji.

Emoticons and emojis can be used for the same purpose. However, emojis have seen a recent surge in popularity, presumably due to the fact that emojis provide colorful graphical representations as well as a richer selection of symbols. In fact, as table 1 shows, there were a large number of emojis in our corpus. A further advantage is that they may transcend language barriers.

We believe that emoticons and emojis can help with assessing the tone of a tweet. Tweets we are interested in will most likely have a negative tone as they reflect people expressing that they are unwell or suffer some symptoms. This means they may contain one or more emojis/emoticons denoting sadness, anger or tiredness, for example. On the other hand the presence of emojis/emoticons denoting happiness and laughter in a tweet may be an indication that the tweet is not relevant to our context of syndromic surveillance.

We investigate also more complex features derived from our words or additional tokens.

#### 3.1.5 Feature construction

##### 3.1.5.1 Word Classes

Following from work by Bergsma et al. [27], we also extended our feature set with further syntactical features in order to make up for the shortcomings words may present when applied to Twitter data. Word classes are labels that Lamb et al. [8] found useful in the context of analysing tweets to categorize them as related to infection or awareness. The idea is that many words can behave similarly with regard to a class label. A list of words is created for different categories such as “*possessive words*” or “*infection words*”. Word classes function similarly to bag of word features in that the presence of a word from a word class in a tweet triggers a count based feature. We manually curated a list of words and classes which are shown in table 4. As we applied lemmatisation and stemming, we did not include multiple inflections of the words in our word classes.

**Table 4.** Our list of word classes with their member words

| Word Class | Member Words |
| --- | --- |
| Infection | sick, down, ill, infect, caught, recover |
| Possession | have, contain, contaminated, my |
| Concern | awful, worried, scared, afraid, terrified, fear, sad, unhappy, feel |
| Humour | laugh, ha, haha, hahaha, lol, lmao, rofl, funny, hilarious, amused |
| Symptomatic | runny nose, cough, spray, shots, wheezing, mucus, cold |

##### 3.1.5.2 Positive and Negative Word Counts

We constructed two dictionaries of positive and negative words respectively. These dictionaries are shown in the appendix. This feature computes for every tweet, the number of positive words and negative words it contains. Words that do not appear in either of our dictionaries are not counted. The classifier should then infer a matching between ratios of positive to negative counts and tweet relevance.

##### 3.1.5.3 Denotes Laughter

This is a simple binary feature which measures the presence of a token (emoji and/or emoticon) that might suggest laughter or positivity. We manually curated and saved a list of positive emojis/emoticons for this. The usefulness of this feature was augmented by also checking for the presence of a small list of more established and popular internet and slang for laughter or humour such as “lol” or “lmao” which stand for “Laughing Out Loud” and “Laughing My Ass Off” respectively. Table 5 shows this feature’s distribution over the data.

**Table 5.** Distribution of some constructed features and classes across the dataset

| Feature | Value Distribution |  | Class Distribution |  |
| --- | --- | --- | --- | --- |
|  |  |  | Relevant | Not Relevant |
| *Denotes Laughter* | TRUE | 3.9% | 31.8% | 68.2% |
|  | FALSE | 96.3% | 24.2% | 75.8% |
| *Negative Emojis/Emoticons* | TRUE | 5.5% | 74.8% | 25.2% |
|  | FALSE | 94.5% | 21.6% | 78.4% |

##### 3.1.5.4 Negative Emojis/Emoticons

This is similar to the *Denotes Laughter* feature but this time looking at the presence of an emoji or emoticon that can be associated with an illness or the symptoms that it may bring., i.e. negative emotions. We decided to include this feature because we discovered ubiquity of emojis on Twitter and wanted to investigate their potential. Table 5 shows this feature’s distribution over the data. We find that this feature may be the most discriminative of the three. Of the instances with a positive value, a high percentage belong to the “relevant” class and of the instances with a negative value, a high percentage belong to the “not relevant” class.

We experimented with two other features - *Contains Asthma-Verb Conjugate* and *Indicates Personal Asthma Report* but found that they underperformed compared to the other features so we do not report on them. We also constructed features from the tweets collected in the latest time period in order to see how the features generalised across time periods. The distributions of the non-continuous features from the latest time period are shown in table 6.

**Table 6.** Distribution of constructed features and classes across tweets from a different time period 2 years apart from our that of our investigated dataset

| Feature | Value Distribution |  | Class Distribution |  |
| --- | --- | --- | --- | --- |
|  |  |  | Relevant | Not Relevant |
| *Denotes Laughter* | TRUE | 4.0% | 13.9% | 86.1% |
|  | FALSE | 96.0% | 32.5% | 67.5% |
| *Negative Emojis/Emoticons* | TRUE | 14.4% | 41.3% | 58.7% |
|  | FALSE | 85.6% | 30.2% | 69.8% |

For each tweet, we appended all of the above features together to form one feature vector. Each tweet *T*_*i*_ is therefore represented by an *f* dimensional vector, where *f* is a sum of the number of terms, *n*, in the constructed vocabulary, and the dimensionality of our custom features *C* (*Word Classes, Positive and Negative Word Counts, Denotes Laughter* and *Negative Emojis/Emoticons*). This gives us

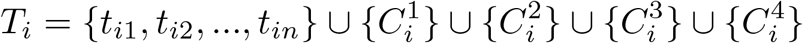

where *t*_*ij*_ represents the weight of the *j*-th vocabulary term in the *i*-th tweet and 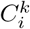 represents the value of the *k*-th custom feature in the *i*-th tweet. The feature vectors are represented in code by dictionary (or hashmap) objects which allows them to contain different types of values (ie. binary, continuous and categorical).

### 3.2 Text Classification

A classification algorithm for text can be used to automatically classify tweets, in this case, to the categories of relevant/not relevant. We first applied a variety of popular and powerful supervised classification algorithms to the data namely - Naive Bayes, Decision Trees, Logistic Regression and Support Vector Machines. We used the Python implementations found in the Natural Language ToolKit (NLTK) and Sci-Kit Learn [28].

Due to the relatively limited number of labelled instances in our data set, we decided to take a semi-supervised approach to learning. We implemented a semi-supervised approach which is suited to small to medium sized datasets [29]. Semi-supervised learning attempts to make use of the combined information from labelled and unlabelled data to exceed the classification performance that would be obtained either by discarding the unlabelled data and applying supervised learning or by discarding the labels and applying unsupervised learning. Our intention is to extend the labelling in a semi-supervised fashion. We make use of the heuristic approach to semi-supervised learning and employ a ***self-training iterative labelling algorithm***. We then extend this work by using a form of ***co-training***.

#### 3.2.1 Self-training model

We adopted an *Iterative Labelling Algorithm* for semi-supervised learning [30]. Iterative labelling algorithms are closely related to and are essentially extensions of the Expectation-Maximization (EM) algorithm put forward by Dempster et al. [31]. The iterative labelling algorithm is a sort of *meta-algorithm* which uses a data set *S* of labelled instances *L*, unlabelled instances *U*, and a supervised learning algorithm *A* with

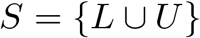

An iterative learning algorithm aims to derive a function *f* which provides a mapping from *S* to a new dataset *S*′:

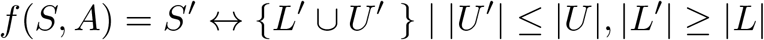

Such an algorithm can be defined simplistically as an iterative execution of three functions: *Choose-Label-Set*(*U, L, A*) selects and returns a new set, *R*, of unlabelled examples to be labelled; *Assign-Labels*(*R, S, A*) generates labels for the instances selected by *Choose-Label-Set*(*U, L, A*); *Stopping-Condition*(*S, S*′) dictates when the algorithm should stop iterating.

##### Algorithm Iterative labelling Algorithm

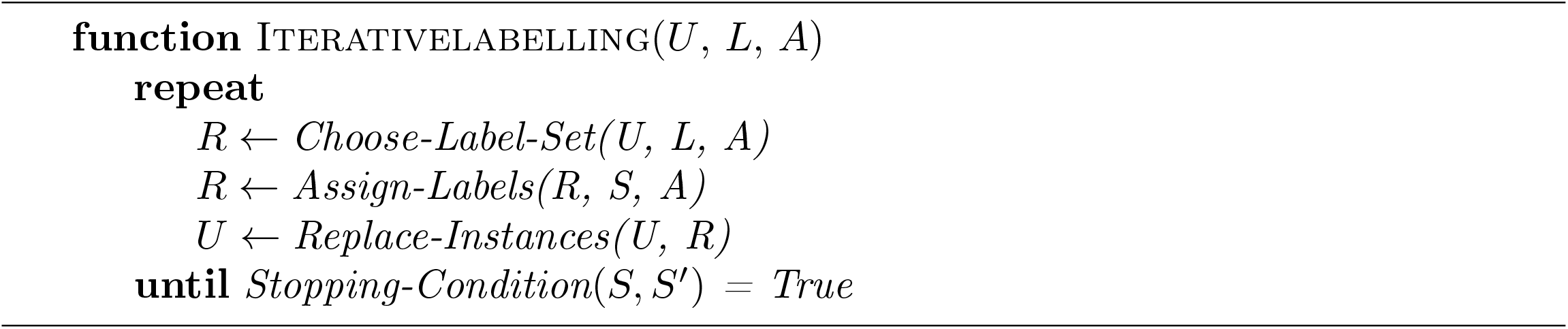

For our choice of supervised learning algorithm, we selected the Logistic Regression classifier after experimenting with different supervised models and finding it to perform best. We used the trained Logistic Regression classifier’s predictions to label unlabelled instances in the *Assign-Labels* function. We set our stopping condition such that the iteration stops when either all the unlabelled data is exhausted or there begins to be a continued deterioration in performance as more data is labelled. Along with the class of an applied instance, we also compute the model’s confidence in its classification. Our algorithm, inspired by Truncated Expectation-Maximization (EM) [32], then grows *L* based on the confidence of our model’s classification. When an instance from *R* is classified, if the confidence of the classification is greater than some set threshold *θ*, the instance is labelled. Considering this, our algorithm falls within the *confidence-based* category of iterative labelling or self-training algorithms because it selects instances for which the trained classifier has a high confidence in its predictions.

Confidence-based iterative labelling algorithms can tend toward excessively conservative updates to the hypothesis, since training on high-confidence examples that the current hypothesis already agrees with will have relatively little effect [32]. In addition, it has been proven that in certain situations, many semi-supervised learning algorithms can significantly degrade the performance relative to strictly supervised learning [33, 34].

#### 3.2.2 Co-training model

To address the problems of self-training, we take some ideas from *co-training* [35] to try to improve our algorithm. Co-training requires different views of the data so that multiple classifiers can be maintained for the purpose of labelling new instances. Recall that each tweet can be represented as a feature vector *T*_*i*_ with various features. We now distinguish two representations. The first is a concatenation of our *Bag-of-Words, Word Classes, Denotes Laughter* and *Negative Emojis/Emoticons* features. We represent this feature space as *X*_1_. The second kind of feature vector is a concatenation of our *Bag-of-Words, Positive and Negative Word Counts, Denotes Laughter* and *Negative Emojis/Emoticons* features. We represent this feature space as *X*_2_. We can think of *X*_1_ as the **taxonomical** feature space as is characterised by its inclusion of the *Word Classes* feature while *X*_2_ can be the **sentimental** feature space and this is characterised by its inclusion of the *Positive and Negative Word Counts* feature. As such, *X*_1_ and *X*_2_ offer different, though overlapping, views of the dataset. Each tweet is then represented as a feature vector from each of these spaces.

We now maintain two separate classifiers trained on different views of the data. During the iterative labelling process, we only label instances for which at least one of the classifiers has a high confidence in its prediction and take the result of that classification as the label. Similar to self-training, at the end of each iteration, the newly labelled data is incorporated into each of the classifiers to update their hypotheses. Once the iterative labelling process is completed, the prior training examples for both classifiers as well as the newly labelled examples are joined together and used to train a new classifier using all the features which will then be applied in practice. The benefit of co-training is that the examples labelled by one classifier are also presented to the other classifier to update the hypothesis on the complementary view. Thus, the examples, as represented in each view, receive at least some of their labels from a source other than the classifier that will be updated with them [30].

#### 3.2.3 Correcting the class imbalance

We started with an initial set of manually labelled data contained 3,500 tweets. This consisted of 24.7% tweets that were labelled as relevant and 76.3% labelled as irrelevant. Imbalanced data causes well known problems to classification models [36]. We initially tried both oversampling and undersampling techniques to create a balanced training dataset. We found no major difference as a result of either technique so opted for under sampling. The class distribution over the balanced training set had 47% of tweets as relevant and 53% as irrelevant. The test set was not balanced.

#### 3.2.4 Performance metrics

Another important aspect of imbalanced data and of classification in general is having the right performance metric for assessment of classification model [37]. Overall accuracy is a misleading measure [38] as it may only be reflecting the prevalence of the majority class. This is called the accuracy paradox, i.e. we could get high accuracy by classifying all tweets as irrelevant. That would, however, not improve our signal. The aim of our endeavour is to identify tweets which might suggest an increase of cases for a particular syndrome (asthma/difficulty breathing) for the purpose of syndromic surveillance. Our signal for some syndromes is quite weak as not many cases may occur at a national level and even less may be talked about on Twitter. Because of this, we are very concerned with identifying and keeping instances of the positive class (relevant tweets). We would like to reduce the number of irrelevant tweets but not at the expense of losing the relevant tweets. This means that, for our classifier errors are not of equal cost. Relevant tweets that are classified as irrelevant or False Negative (FN) errors should have a higher cost and hence be minimised; we can have more tolerance of irrelevant tweets classified as relevant or False Positive (FP) errors. Those subtleties are well captured by alternative measures of model performance [39] such as Recall, the probability that a relevant tweet is identified by the model, defined as

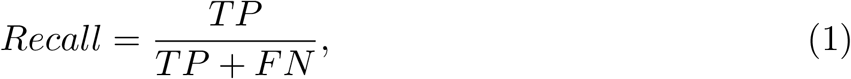

and Precision, the probability that a tweet predicted as relevant is actually relevant

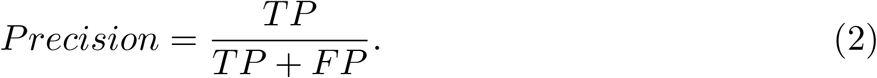

where TP and TN stand for True Positives and True Negatives respectively.

Precision and recall are often trading quantities. A measure that combines precision and recall is the *F*-measure or *F*-score [40]. The generic version of this is:

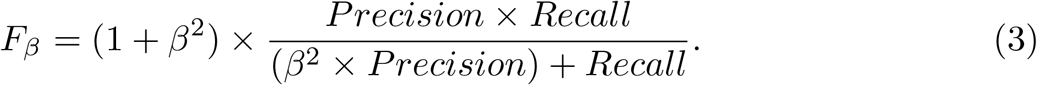

With *β* = 1, that becomes the traditional or balanced *F*_1_-measure. With a *β* = 2, the *F*_2_ measure weighs recall higher than precision and so it may be more suited to our purpose.

#### 3.2.5 Assessment of features and key words

We also assessed the discriminative ability of each of our features by performing feature ablation experiments [41]. We evaluated the performance of a given classifier when using all our features, and then again after removing each one of these features. The difference in the performance is used as a measure of the importance of the feature. We chose to use the difference in *F*_1_ metric over *F*_2_ in this analysis because we wanted to convey how the features performed in the general task of tweet classification.

We also performed some analysis on the word features to learn which words in our vocabulary were the best indicators of relevant tweets. We analysed the bag-of-words component of our compound feature vectors in order to calculate the *informativeness*, or *information gain* of each word unigram. The information gain of each feature pair is based on the prior probability of the feature pair occurring for each class label. A higher information gain (hence, a more informative feature,) is a feature which occurs primarily in one class and not in the other. Similarly, less informative features are features which occur evenly in both classes. The information gain idea is pivotal to the decision tree algorithm but generalizes to others and was adapted in the NLTK package for use in a broader sense. In NLTK, informativeness of a word *w* was calculated as the highest value of *P* (*w* = *feature*_*value|class*) for any class, divided by the lowest value of *P* (*w* = *feature*_*value|class*) [28]. This informativeness *I*, is summarised below:

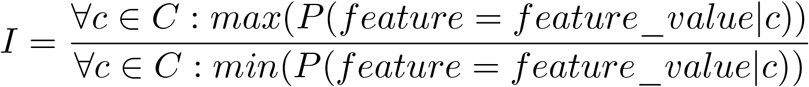

where *C* is the set of all classes and *c* is a possible class.

Recall that to collect tweets, we made use of Twitter’s streaming API which allowed us to specify keywords to restrict the data collection to tweets containing those specific terms. We measured the usefulness of the keywords we selected. To do this, we assessed their information retrieval performance. Specifically, we used the precision-recall metric. In an information retrieval context, precision and recall are defined in terms of a set of retrieved documents and their relevance. We use our original set of labelled tweets for this assessment (i.e. the set of 3500 tweets). In our scenario, the labelled tweets make up the set of retrieved documents and the tweets labelled as belonging to the “relevant” class make up the set of relevant documents. In this context, recall measures the fraction of relevant tweets that are successfully retrieved while precision measures the fraction of retrieved tweets that are relevant to the query.

## 4 Results

### 4.1 Classifiers

The results of our fully supervised and semi-supervised classification are presented in table 7. The original data was divided into 70:30 training and test split through random sampling and the results presented are measures obtained from the test data. Of the fully supervised classifiers, Logistic Regression and SVM are very sensitive to hyperparameters. In assessing the performance of these classifiers, we varied and tuned the hyperparameters until we found the best results and reported those. For Logistic regression, we used L2 regularization with a regularization strength *C* of 0.00001. For the SVM, we used a Radial Basis Function kernel and *C* of 0.01. For the iterative labelling experiments, we varied and tuned the confidence thresholds until we found the best results and reported those. Below, we also discuss in more detail how the confidence threshold affected the iterative labelling performance as it is a key aspect of the algorithms. The best fully supervised approach according to a combination of the *F*_1_ and *F*_2_ scores was the Logistic Regression classifier, which achieved an *F*_2_ score of **0.764** on the test data. This equated to an overall prediction accuracy of **91.5%**. The best semi-supervised approach, which was the co-training algorithm, achieved an *F*_2_ score of 0.903 on the test data, with an overall predictive accuracy of **92.3%**. Overall, the semi-supervised approach is more accurate and achieves higher *F* scores. To confirm what we concluded from the results, we applied a paired *t*-test to test the difference in *F*_2_ scores between the fully supervised logistic regression algorithm and the co-training algorithm. Before carrying out this test, we confirmed that the data satisfied the assumptions necessary for the paired t-test to be relevant - continuous, independent, normally distributed data without outliers. This resulted in a *t*-statistic of 18 and a **p-value of 9.6 × 10**^**−14**^ which suggests that the difference between the *F*_2_ scores of the two algorithms was not due to chance.

**Table 7.**
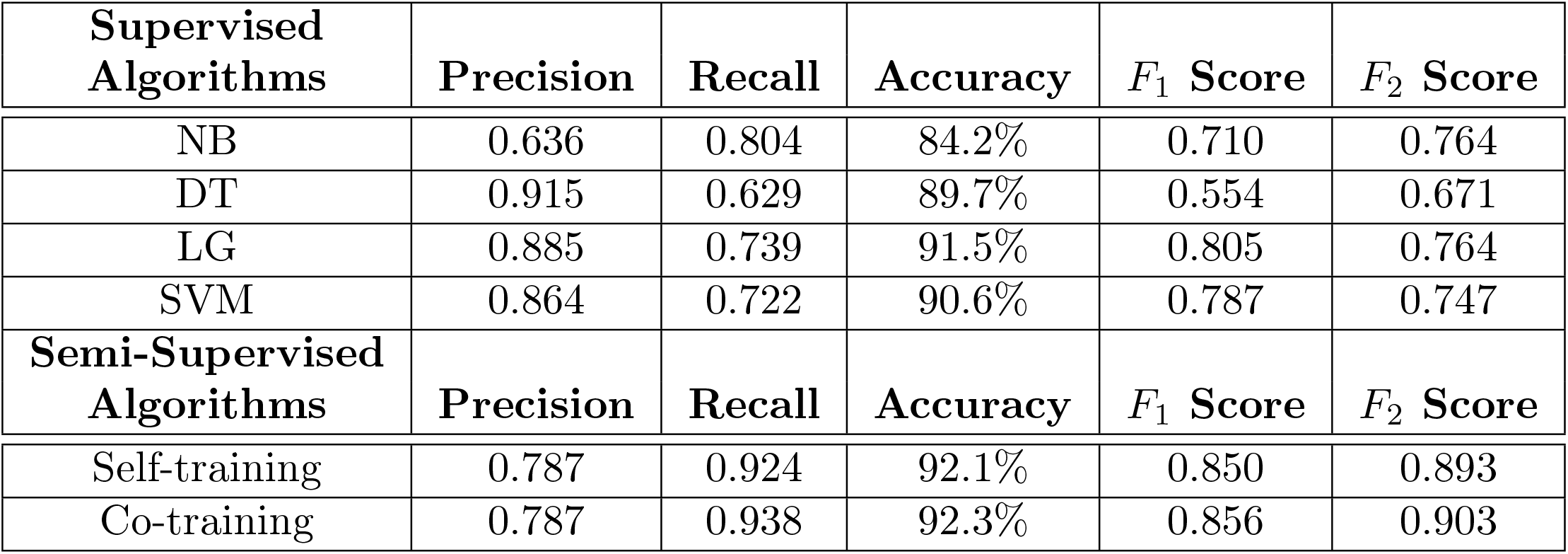
Results of relevance classification on the test data. Naive Bayes (NB), Decision Tree (DT), Logistic Regression (LR) and Support Vector Machine (SVM) algorithms are reported together with the self-training and co-training iterative labelling algorithms.

| Supervised Algorithms | Precision | Recall | Accuracy | $F_1$ Score | $F_2$ Score |
| --- | --- | --- | --- | --- | --- |
| NB | 0.636 | 0.804 | 84.2% | 0.710 | 0.764 |
| DT | 0.915 | 0.629 | 89.7% | 0.554 | 0.671 |
| LG | 0.885 | 0.739 | 91.5% | 0.805 | 0.764 |
| SVM | 0.864 | 0.722 | 90.6% | 0.787 | 0.747 |
| Semi-Supervised Algorithms | Precision | Recall | Accuracy | $F_1$ Score | $F_2$ Score |
| Self-training | 0.787 | 0.924 | 92.1% | 0.850 | 0.893 |
| Co-training | 0.787 | 0.938 | 92.3% | 0.856 | 0.903 |

To give a better understanding of how the different measures manage to balance the number of FP and FN. We also present the confusion matrices for both the best performing fully supervised and semi-supervised methods on the test data. These confusion matrices are shown in tables 8 and 9 respectively. We see that the supervised approach shown in Table 8 will retain 243 tweets in total (predicted as positive). Of those 215 are positive tweets retained, however 76 positive tweets will be discarded. In contrast, the semi supervised approach shown in Table 6 will retain 347 tweets, of those 273 will be positive, with only 18 positive tweets being discarded. From the confusion matrices, we see that the semi-supervised approach performs better for the purpose of syndromic surveillance as it yields only 18 false negatives even though it also yields 74 false positives. Considering that our aim is to develop a filtering system to identify the few relevant tweets in order to register a signal for syndromic surveillance it is critical to have high recall, hopefully accompanied by high Precision, and therefore high accuracy. The semi-supervised method is able to identify and retain relevant tweets more often, while also being able to identify irrelevant tweets to a reasonable degree. Hence even with a shortage of labelled data the semi-supervised algorithms can be used to filter and retain relevant tweets effectively.

**Table 8.**
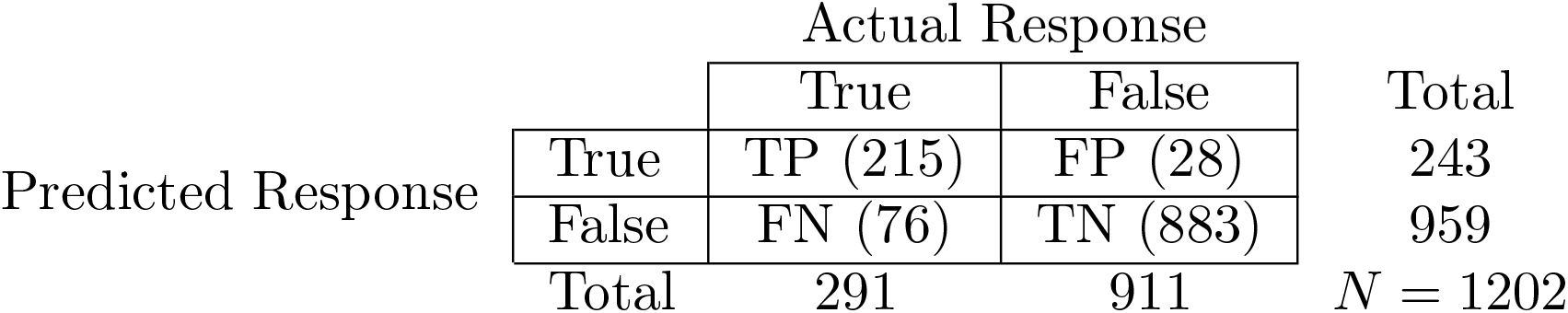
Confusion matrix for Logistic Regression fully supervised classification on the test data

**Table 9.**
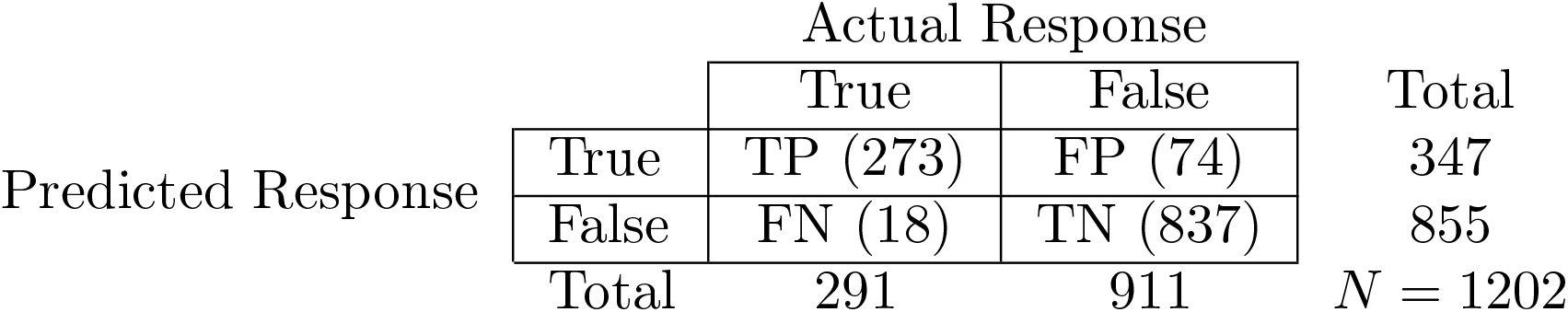
Confusion matrix for Co-training semi-supervised algorithm on the test data

Fig 2 shows how the performances of the semi-supervised systems change as the confidence threshold changes. The confidence threshold controls how conservatively the semi-supervised system assimilates unlabelled instances as it represents how confident the semi-supervised system needs to be in its classification before assimilating the instance to inform future decisions. We observed co-training with logistic regression to perform best. We also observed that for lower confidence thresholds between 0.1 and 0.5, self-training performance is usually lower and does not change much between thresholds. Co-training on the other hand, appears to be less sensitive this parameter. Fig 2 also reiterates what we learned from table 7 that logistic regression is our strongest fully supervised model and shows us that it is the most robust to different confidence thresholds when used in an iterative labelling context.

**Fig 2.**
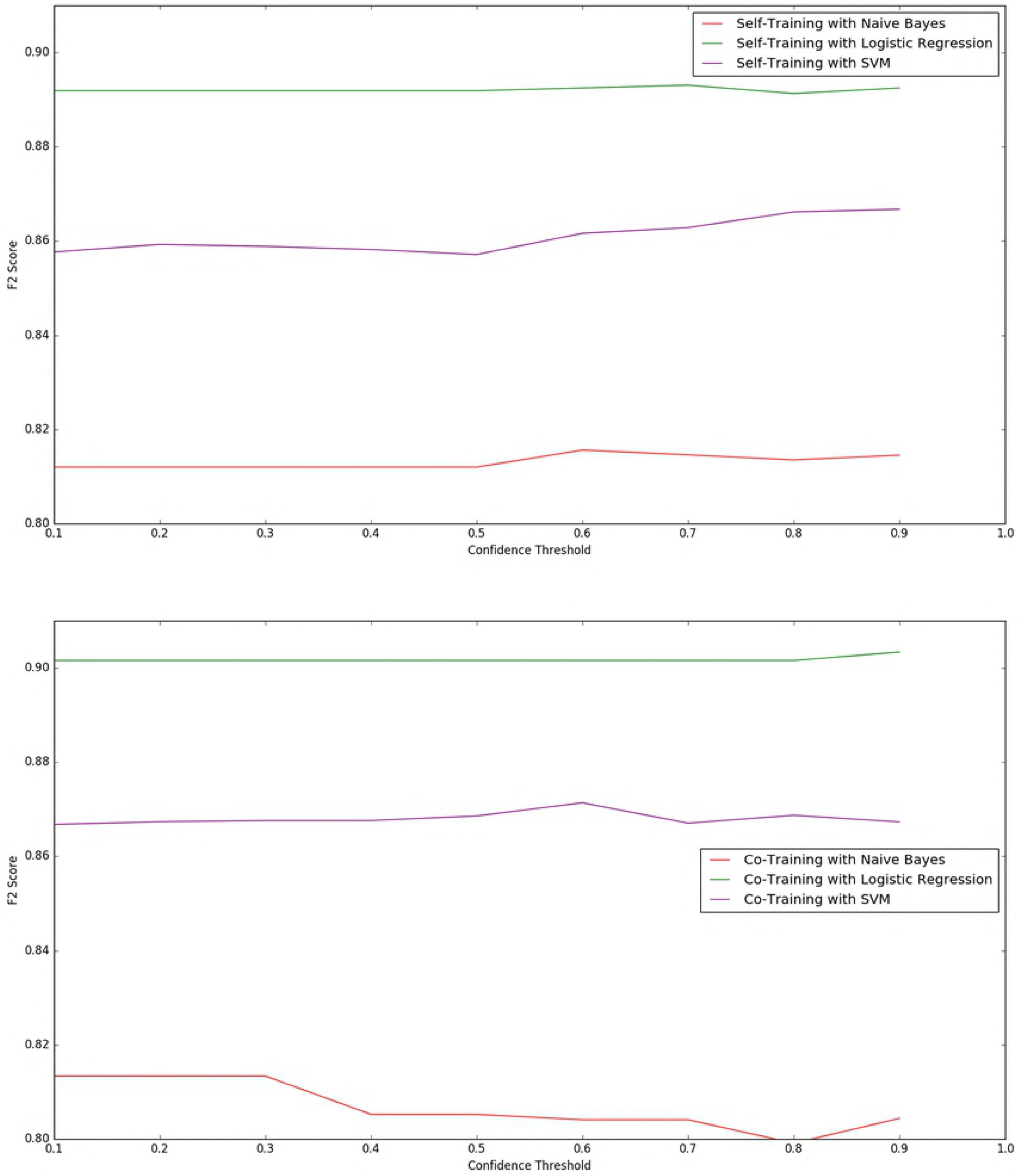
Graph of F2 performance of Iterative Labelling using different confidence thresholds

The main issue with iterative labelling algorithms is that, because the classifiers are not perfect and do not have 100% accuracy, we cannot be sure that the unlabelled instances that they label for assimilation are always correct. This means that how well they initially perform before starting any iterations is vital. Consider a classifier, initially of poor performance (with an accuracy of 0.2 for example). When classifying unlabelled instance with which to train itself, 80% of its classifications will be wrong, so it will assimilate false hypotheses, which will in turn make its performance in the next iteration even worse and so on. Conversely, if the initial accuracy is high, it is more likely to correctly classify unlabelled instance and be less resistant to the drop in performance from assimilating false hypotheses. We conducted an experiment to measure the quality of the automatically labelled instances assimilated by our semi-supervised classifiers. For this exercise, we used the second set of labelled tweets from a different time period as the “unlabelled” set with to which the iterative labelling is applied to. The same training set as in our other experiments was used for the initial training stage. The self-training and co-training processes were initiated, applying these classifiers to the alternative set of labelled data (around 2000 instances) in steps of 200. Fig 3 shows a plot of the proportion of correctly classified instances that the iterative labelling process assimilated. The co-training approach had a higher rate of being correct when making new additions. This was in fact the aim of adopting co-training with its multiple different views of the same data. The proportion of correct assimilations of both the self-training and co-training methods rises as more data is assimilated, due to the fact that the systems are getting more intelligent. Although we could not test beyond 2000 instances (because of our limited labelled data), we believe that the proportion of correct assimilations will increase until a certain point, after which it will plateau. We expect this plateau due to the fact that at a certain point, the iterative learning classifiers will have nothing new to learn from new data after having been exposed to so much.

**Fig 3.**
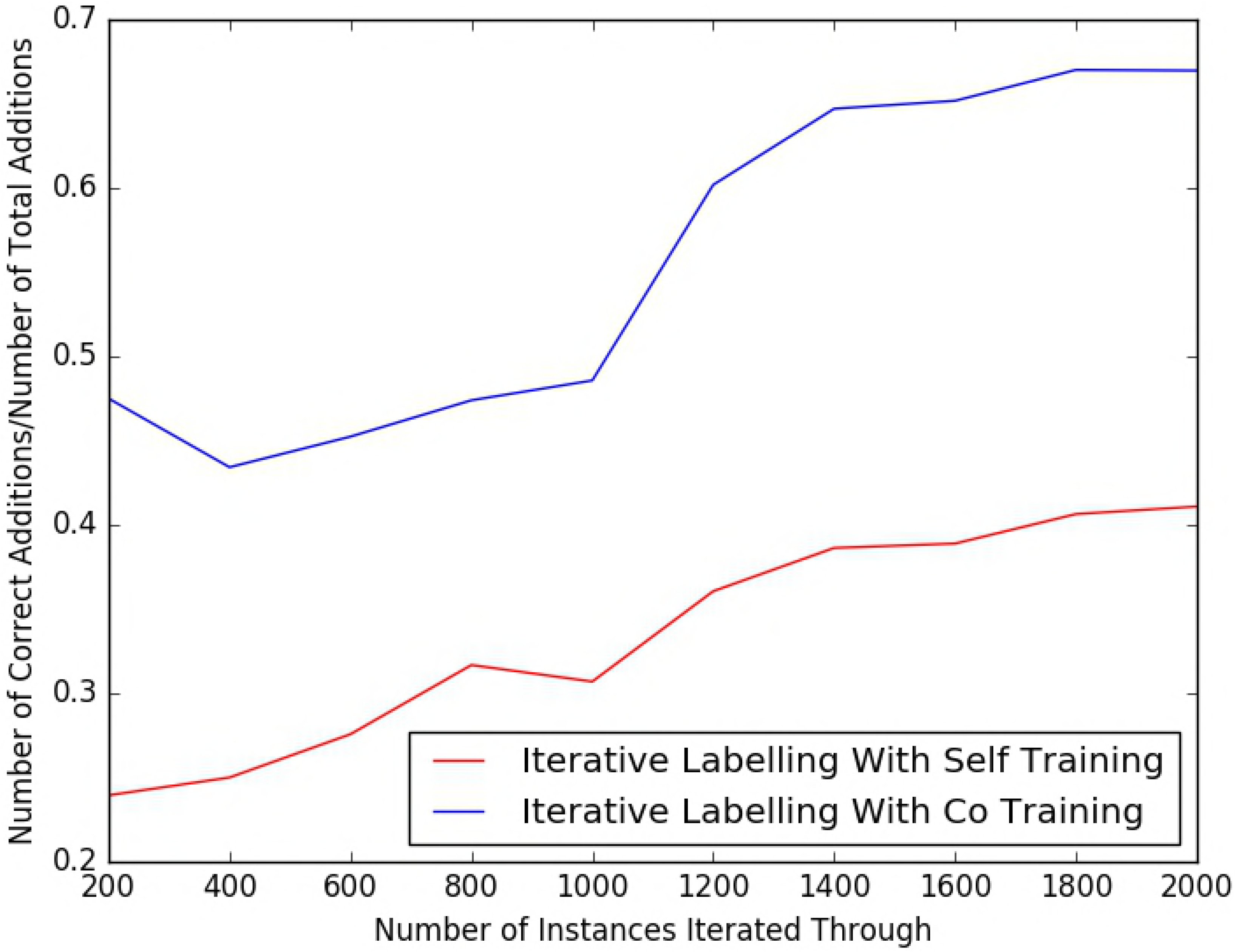
Graph showing how many correct assimilations the iterative labeling algorithms make per iteration using labelled data from a different time period

As with the features constructed, we tested how the classifiers would perform for new data collected at a different time period to assess if shifts in language and colloquialisms could have an impact on performance. Our classifiers were built on data from the first collection period (see Table 2). For a simple assessment, we applied our trained model to tweets collected in the most recent collection period, which had a time gap of two years from the original data. Our semi-supervised approach based on co-training achieved a precision of 0.400 and a recall of 0.628 on the 2,000 labelled tweets from the most recent collection period. This means an *F*_1_ score of 0.488 and more importantly, an *F*_2_ score of 0.564. For comparison purposes, we also applied the fully supervised logistic regression algorithm to the data from this new time period. This yielded a precision of 0.510 and a recall of 0.419. This meant an *F*_1_ score of 0.460 and an *F*_2_ score of 0.434. In both cases, we observe a deterioration in performance when introduced to tweets from a different time period. This poses an important issue to consider about how language online changes moving forward. Although it changes very gradually, after a period of one or two years, the changes are substantial enough to render the natural language-based models less effective.

### 4.2 Feature Analysis

Table 10 shows the results of the feature ablation experiments. We found that negative emojis/emoticons were the most discriminative of our features followed by the *Denotes Laughter* feature in the supervised approach, which also captures emojis as well as colloquialisms, and *Positive/Negative Word Count* in the semi-supervised approach. All three of these features capture the mood of a tweet. We also found that our additional features proved effective for classifying tweets as relevant or not and yielded an improvement over the bag-of-words baseline.

**Table 10.** F1 scores after feature ablation

| Ablated Feature | Supervised $F_1$ Score | Semi-supervised $F_1$ Score |
| --- | --- | --- |
| *None* | 0.710 | 0.714 |
| *Denotes Laughter* | 0.628 | 0.643 |
| *Negative Emojis/Emoticons* | **0.627** | **0.620** |
| *Word Classes* | 0.677 | 0.637 |
| *Positive/Negative Word Count* | 0.648 | 0.625 |

Table 11 shows the words found to be most informative. For example, the table shows that, of the tweets containing the word *chest*, 96% are relevant and only 4% are irrelevant. The training data is used for this calculation. A surprising negative predictor was the word *health*. When *health* appeared in a tweet, the tweet was irrelevant 94% of the time. The word *pollution* shows a similar trend. This suggests that when Twitter users are expressing health issues, they may not use precise or formal terms, opting for simple symptomatic and emotional words such as *chest, cold* or *wow*. The more formal terms may be more often associated with news items or general chat or discussion. Using this information, we could include some of the more relevant but perhaps unexpected keywords as keywords when collecting streaming tweets from Twitter in order to better target and collect relevant tweets.

**Table 11.** Most informative words measured by their *Informativeness* and their relevant to irrelevant prior probabilities

| Word | $I$ (Relevant:Irrelevant) | Relevant Prior Probability | Irrelevant Prior Probability |
| --- | --- | --- | --- |
| chest | 22/4 | 0.96 | 0.04 |
| throat | 17/1 | 0.95 | 0.05 |
| wow | 17/1 | 0.95 | 0.05 |
| health | 1/17 | 0.06 | 0.94 |
| cold | 16/1 | 0.94 | 0.06 |
| moment | 15/1 | 0.94 | 0.06 |
| forecast | 1/14 | 0.07 | 0.93 |
| awake | 13/1 | 0.93 | 0.07 |
| awful | 13/1 | 0.93 | 0.07 |
| sick | 13/1 | 0.93 | 0.07 |
| cough | 12/1 | 0.92 | 0.08 |
| pollution | 1/12 | 0.08 | 0.92 |
| bed | 11/1 | 0.91 | 0.09 |
| hate | 11/1 | 0.91 | 0.09 |
| watch | 10/1 | 0.91 | 0.09 |

We also investigated which emojis where most prevalent in our data set as well as how often each emoji showed up in a tweet of each class. Fig 4 shows the frequency with which each emoji occurred in the labelled tweets. It shows that only a few emojis appear very frequently in tweets collected in our context. This means that only a few important emojis were needed for determining tweet relevancy as opposed to monitoring for the full emoji dictionary. Table 12 shows a list of some emojis and the distribution of classes that tweets belonged to whenever they contained said emoji. Overall, it can be seen that each of these emojis tend to lean heavily towards one class. This shows that they could be quite discriminative and useful indicators of class membership hence helpful features.

**Table 12.**
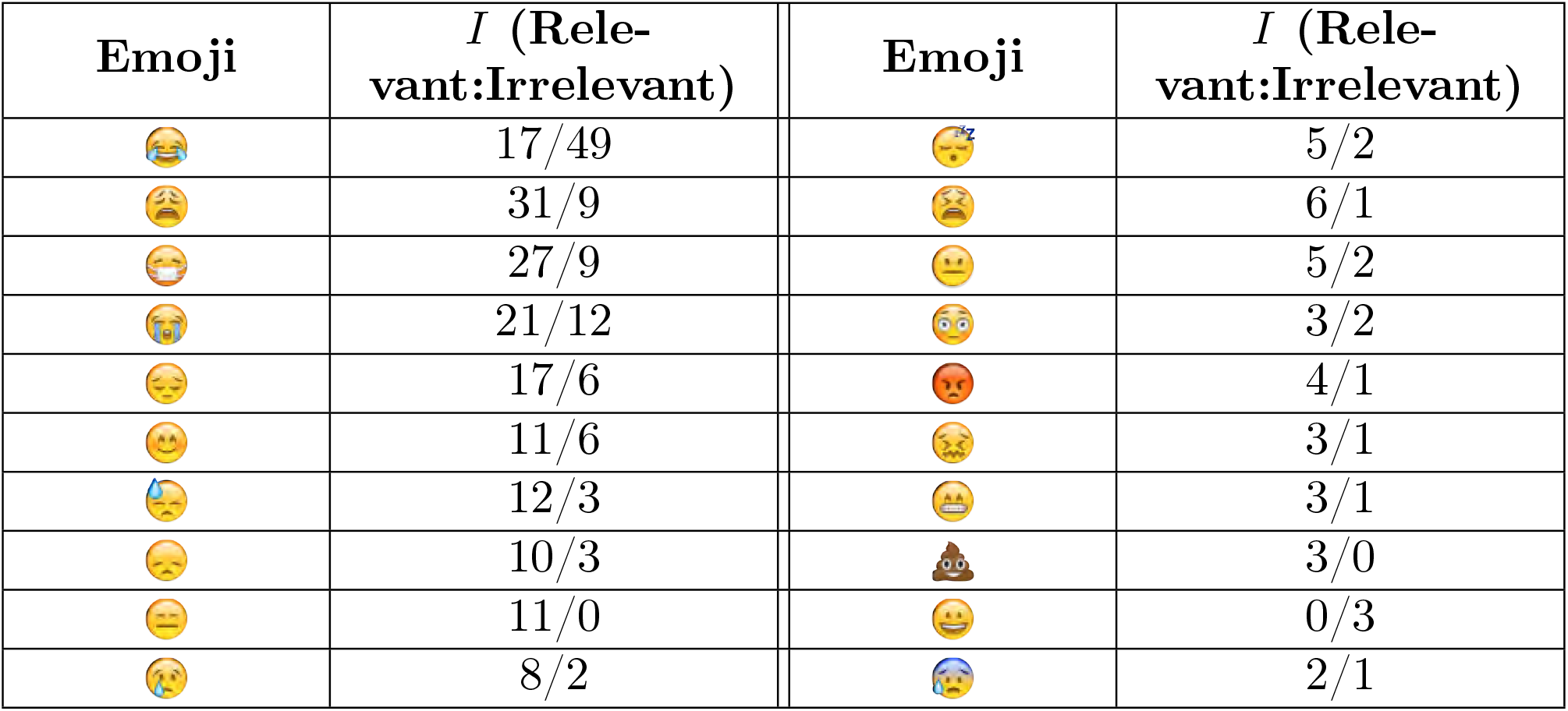
Most frequent emojis in labelled data and their distributions

**Fig 4.**
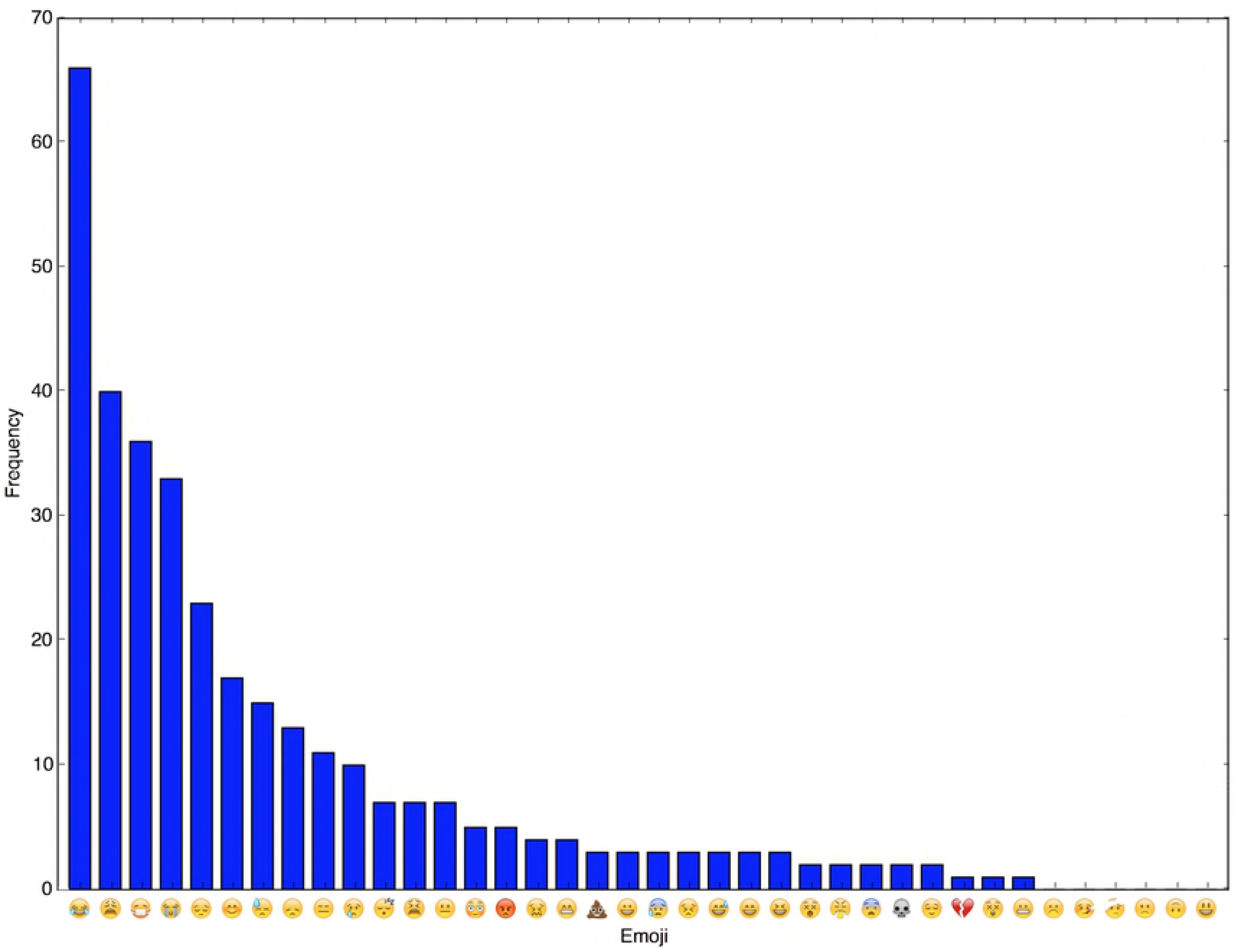
Bar chart showing emoji frequency in labelled data

### 4.3 Keyword Analysis

Table 13 shows the results of the assessment of key words used in tweet collection. We found that *asthma, pollution* and *air pollution* were the keywords that yielded the most results at 1313, 757 and 509 out of a total of 3500. *Wheezing, fumes* and *inhaler* were next with 219, 132, 121 tweets respectively. The remaining keywords return very few results (below 40) or no results. *Asthma* had the highest recall but not very high precision so most of its results were irrelevant. *Wheezing, inhaler, wheeze, cannot breathe, can’t breathe, difficulty breathing* and *short of breath* have good precision although their recall is not that high. Some of those keywords express direct symptoms of the syndrome under investigation, hence, we expect good precision. *Tight chest* and *pea souper* have very high precision but only appeared in two tweets each. Of the keywords used, *wheezing* was the most useful in that it brought in a lot of results, most of which were relevant. We included a common misspelling of the keyword with the highest recall power - *asma*. We found that *asma* only appeared in 4 tweets. We hypothesize that this is due to the fact that most users of Twitter post from devices capable of autocorrect hence it may not be necessary to worry about misspelling of keywords.

**Table 13.** Assessment of the retrieval quality of the search keywords

| Keyword | Precision | Recall | Keyword | Precision | Recall |
| --- | --- | --- | --- | --- | --- |
| asthma | 0.174 | 0.475 | poor air quality | 0.000 | 0.000 |
| pollution | 0.009 | 0.015 | murk | 0.000 | 0.000 |
| air pollution | 0.008 | 0.008 | can't breathe | 0.556 | 0.010 |
| wheezing | 0.406 | 0.185 | difficulty breathing | 0.125 | 0.002 |
| fumes | 0.030 | 0.008 | short of breath | 0.333 | 0.004 |
| inhaler | 0.198 | 0.050 | respiratory disease | 0.000 | 0.000 |
| smog | 0.023 | 0.002 | asma | 0.000 | 0.000 |
| gasping | 0.025 | 0.002 | tight chest | 0.500 | 0.002 |
| puffing | 0.033 | 0.002 | pea souper | 0.500 | 0.002 |
| wheeze | 0.138 | 0.008 | itchy eyes | 0.000 | 0.000 |
| panting | 0.043 | 0.002 | could not breathe | 0.000 | 0.000 |
| cannot breathe | 0.412 | 0.015 | couldn't breathe | 0.000 | 0.000 |
| trouble breathing | 0.100 | 0.002 | chest tightness | 0.000 | 0.000 |
| sore eyes | 0.100 | 0.002 | acid rain | 0.000 | 0.000 |

The informativeness, *I*, was calculated when the keywords were also features in the classifiers and is presented in Table 14. Most of the keywords were not informative as features with an informativeness ratio of 1:1 for relevant:irrelevant tweets so they are not included. We found some overlap where streaming keywords where informative in the relevance model though not always associated with the relevant class. For example, *Pollution*, which was a keyword, appeared in the ranking of top 15 most informative words though associated with the irrelevant class. Similarly *fumes* also associates with the irrelevant class more. On the other hand *wheezing* had good information retrieval performance and associated with relevant tweets.

**Table 14.** Keyword Informativeness *I* of keywords

| Streaming Keyword | $I$ (Relevant:Irrelevant) | Relevant Prior Probability | Irrelevant Prior Probability |
| --- | --- | --- | --- |
| pollution | 1/12 | 0.08 | 0.92 |
| wheezing | 5/1 | 0.84 | 0.16 |
| fumes | 1/3 | 0.24 | 0.76 |
| panting | 1.6/1.0 | 0.62 | 0.38 |

### 4.4 Syndromic Surveillance Comparisons

As we discussed earlier, the purpose of our relevance filtering is a syndromic surveillance application. After constructing and evaluating our semi-supervised filtering systems, we assessed their utility for syndromic surveillance purposes by retrospectively applying them to Twitter data and comparing the results against data for existing syndromic indicators from the Public Health England (PHE) Real-time Syndromic Surveillance Service. For this experiment, we made use of unlabelled Tweets from our second collection period, June to August 2016. Syndromic surveillance data on the proportion of daily GP Out-of-Hours (GPOOH) calls for ***asthma, wheezing or difficulty breathing*** and telehealth calls (NHS 111) for ***difficulty breathing*** for the period June to August 2016 were compared to the signal detected from our Twitter data using our semi-supervised and supervised filtering systems. As a sense check, we also compared our detected Twitter signal time series against non-respiratory syndrome data in the form of NHS 111 calls for ***diarrhoea***. This was to provide some form of control which should not correlate with the Twitter asthma signal.

The resulting time series shows the daily proportion of relevant symptomatic tweets and consultations/calls as observed on Twitter and recorded by PHE (Fig 5 and Fig 6). The signals were smoothed using a 7-day moving average to remove the fluctuations in daily activity for GPOOH data as that service receives more usage over the weekends. We also included a time series showing the Twitter signal without any filtering for further perspective. We see that the time series plots of the self-training and co-training filtering follow a similar trend to the GP data time series. Also, the time series for the Twitter data without any filtering has lots of spurious peaks in relation to the ground truth data (i.e. the syndromic surveillance data). Both of these observations together suggest that Twitter data might mirror the health activity of a population and that relevance filtering is useful in reducing noise and obtaining a clearer picture of such activity. Additionally, we see that while the unfiltered Twitter signal does not match well with the *asthma/wheeze/difficulty breathing* or *difficulty breathing* signal, it still seems to match better than that of the diarrhoea signal.

**Fig 5. Time series plots comparing GP asthma/wheeze/difficulty breathing data to signals from supervised and semi-supervised Twitter analysis and unrelated diarrhoea.**

**Fig 6. Time series plots comparing NHS 111 difficulty breathing data to signals from supervised and semi-supervised Twitter analysis and unrelated diarrhoea.**

To further evaluate the quality of our detected signal, we calculated a Pearson correlation coefficient to determine the strength and direction of any monotonic relationship between the indicators (table 15). We observed a weak but statistically significant correlation between the Twitter signals and the asthma and difficulty breathing syndromic surveillance data. For the diarrhoea syndrome, there was no statistically significant correlation with the Twitter time series. To corroborate this, it can be seen from the time series that the plots for our detected Twitter signals and GP data follow a similar downward trend but the diarrhoea signal does not. This further suggests that our semi-supervised system can detect activity similar to that detected by traditional syndromic surveillance systems recording GP visits and NHS 111 calls. This suggests that they could potentially be further explored as an additional form of syndromic surveillance.

**Table 15.** Pearson correlations and P-Values for detected signals with syndromic surveillance signals

|  | Self-Training | Co-Training | Logistic Regression |
| --- | --- | --- | --- |
| Asthma/Wheezing/DB | 0.249($p = 0.04$) | 0.414($p = 0.0004$) | 0.255($p = 0.03$) |
| Difficulty Breathing | 0.228($p = 0.04$) | 0.424($p = 0.0002$) | 0.214($p = 0.07$) |
| Diarrhoea | -0.01($p = 0.9$) | 0.04($p = 0.7$) | 0.05($p = 0.7$) |

## 5 Discussion

Twitter is a noisy data source for syndromic surveillance but through data processing and tweet classification, we have been able to identify relevant tweets among the noisy data for a specific syndrome or incident (asthma/difficulty breathing). This in turn allowed us to extract a signal of potential use for syndromic surveillance that correlated positively with real-world public health data. Using a semi-supervised method of classification for filtering tweets, we achieve an accuracy of 92.3% and *F*_1_ and *F*_2_ scores of 0.856 and 0.903 respectively. We argued that recall is very important for us because we want to keep all the relevant tweets so that we can have some signal, even if amplified by some misclassified irrelevant tweets. The best recall, obtained by the semi-supervised algorithm equated to retaining over 90% of the relevant tweets after classification. Also, the semi-supervised approach allowed us to use 8000 previously unlabelled tweets before it started to see a deterioration in performance. This allowed us to make use of data that is very easily collected without it going to waste.

Tweet classification using supervised learning has received a lot of attention [14, 15, 17, 42, 43] and gave us good results with *F*_1_ and *F*_2_ scores of 0.805 and 0.764 respectively for the Logistic Regression method. Tweet labelling, required for supervised classification, is time consuming, however, so often researchers do not use all of the data available to build the model. Semi-supervised methods for tweet classification have been used for sentiment analysis [44, 45]. They can enable more of the collected data to be used for training the classifier bypassing some of the labelling effort. Johnson el al. [46] used a method called label propagation and reported accuracy of 78%. Baugh [20] proposed a hierarchical classification system with self-training and reported accuracy of 61% and an *F*_1_ score of 0.54. We have implemented an iterative labelling semi-supervised approach which seems to have competitive performance and also enables us to use more of the training data without the effort of labelling. Furthermore, we get an improvement on recall over the supervised method, which is important given that the signal we are trying to preserve for syndromic surveillance may be weak. We compare our semi-supervised system to others above but we acknowledge that applications in different domains might weaken the comparison. Baugh [20] also applied semi-supervised systems to tweet classification but not for syndromic surveillance so this comparison might be of more value.

We have also identified strong and novel features in the context of tweet classification: emojis. We have hinted at the growing use of emojis [47] and their importance in establishing the tone of at tweet which in turn is important to relevance classification. Emojis cross language boundaries and are often used by people expressing conditions of interest to syndromic surveillance. Our custom features constructed based on Twitter colloquialisms including emojis proved effective in improving classification performance. Of all our custom features, the one that stood out most was the *Negative Emojis/Emoticons* feature. Emoticons have been used previously [8]. Emojis work even better than emoticons and their uniformity is a real advantage. A smile emoticon could be illustrated in the form “:-D” or “:D”. However, because emojis are actually unicode encoded pictographs with a set standard [48], there exist no variants of the same emoji. In a learning scenario, this reduces fragmentation or duplication of features making them more ideal as features than emoticons.

In terms of geolocation of tweets, we have found that most of the obvious location indicators are not well populated, and those that are, may not be accurate. Hence, future work must tackle geolocation as a real part of the problem for establishing a proper signal from Twitter. After comparing our extracted Twitter signal to real world syndromic surveillance data, we found a positive, albeit weak correlation. This suggests that there is a relationship between asthma related Twitter activity and syndromic surveillance data for asthma and breathing-related incidents. While the actual correlation value indicates a weak relationship, it still suggests that we can detect relevant activity on Twitter which is similar or complementary to that which is collected by traditional means. The strength of the correlation might be affected by the weak location filtering that we have been able to perform. As we discussed, the syndromic surveillance data relates to England but the Twitter data has only been located (not accurately) to the UK. As future work, we plan to assess the full detection capability of Twitter by repeating this analysis prospectively over a longer time period, and for different syndromes, allowing us to determine whether Twitter can detect activity that is of potential benefit to syndromic surveillance.

We also found that “what to collect” is problematic as the data collection of tweets by keywords requires a carefully chosen list of keywords. Furthermore, our experimentation with different type of features like emojis also tell us that the vocabulary used in Twitter is different to expression in other settings (e.g. as part of a medical consultation). Hence we may need to widen our data collection terms to include emojis, emoticons and other types of informal expressions. We may also need to develop adaptive systems in which the set of data collection keywords is dynamically updated to collect truly relevant tweets. So an idea for future research is to begin with a set of keywords, collect tweets, perform relevance analysis and then update the keyword/token list to reflect those that associate with the most relevant tweets, eliminating any keywords/tokens that are not performing adequately.

We saw also that vocabulary and use of tokens change over time. *Negative emojis/emoticons* appeared more often in the second time period, up from 5.5 % to 14.4% of labelled tweets containing them. This could suggest that over the past two years, the use of emojis as a form of expression has grown. However their prevalence in each class also changed, which may explain the classification performance showing some marked deterioration in precision. We performed our research on data collected within a two year period, but further data collection and experimentation would be beneficial to understand the temporality of models generated as Twitter conversations change over time.

## Supporting information

**S1 List Appendix. Twitter data collection keywords** pollution, smog, poor air quality, wheeze, wheezing, difficulty breathing, asthma, inhaler, air pollution, itchy eyes, sore eyes, trouble breathing, cannot breathe, could not breathe, can’t breathe, coudn’t breathe, asma, short of breath, tight chest, chest tightness, respiratory disease, pea souper, murk, fumes, acid rain, gasping, puffing, panting.

**S2 List Appendix. Positive Word Dictionary** adore, adorable, accomplish, achievement, achieve, action, active, admire, adventure, agree, agreeable, amaze, amazing, angel, approve, attractive, awesome, beautiful, brilliant, bubbly, calm, celebrate, celebrating, charming, cheery, cheer, clean, congratulation, cool, cute, divine, earnest, easy, ecstasy, ecstatic, effective, effective, efficient, effortless, elegant, enchanting, encouraging, energetic, energized, enthusiastic, enthusiasm, excellent, exciting, excited, fabulous, fair, familiar, famous, fantastic, fine, fit, fortunate, free, fresh, friend, fun, generous, genius, glowing, good, great, grin, handsome, happy, hilarious, hilarity, lmao, lol, rofl, haha, healthy, ideal, impressive, independent, intellectual, intelligent, inventive, joy, keen, laugh, legendary, light, lively, lovely, lucky, marvel, nice, okay, paradise, perfect, pleasant, popular, positive, powerful, pretty, progress, proud, quality, refresh, restore, right, smile, success, sunny, super, wealthy, money, cash, well, wonderful, wow, yes, yum

**S3 List Appendix. Negative Word Dictionary** abysmal, adverse, alarming, angry, rage, annoy, anxious, anxiety, attack, appalling, atrocious, awful, bad, broken, can’t, not, cant, cannot, cold, collapse, crazy, cruel, cry, damage, damaging, depressed, depression, dirty, disease, disgust, distress, don’t, dont, dreading, dreadful, dreary, fail, fear, scare, feeble, foul, fright, ghastly, grave, greed, grim, gross, grotesque, gruesome, guilty, hard, harm, hate, hideous, horrible, hostile, hurt, icky, ill, impossible, injure, injury, jealous, lose, lousy, messy, nasty, negative, never, no, nonsense, crap, shit, fuck, fukk, fuxk, nausea, nauseous, pain, reject, repulsive, repulse, revenge, revolting, rotten, rude, ruthless, sad, scary, severe, sick, slimy, smelly, sorry, sticky, stinky, stormy, stress, stuck, stupid, tense, terrible, terrifying, threaten, ugly, unfair, unhappy, unhealthy, unjust, unlucky, unpleasant, upset, unwanted, unwelcome, vile, wary, weary, wicked, worthless, wound, yell, yucky

## Acknowledgments

We acknowledge support from NHS 111 and NHS Digital for their assistance and support with the NHS 111 system; Out-of-Hours providers submitting data to the GPOOH syndromic surveillance and Advanced Heath & Care. The authors also acknowledge support from the Public Health England Real-time Syndromic Surveillance Team. Beatriz De La Iglesia and Iain Lake receive support from the National Institute for Health Research Health Protection Research Unit (NIHR HPRU) in Emergency Preparedness and Response.

